# The cervicovaginal microbiome of pregnant people living with HIV on antiretroviral therapy in the Democratic Republic of Congo: A Pilot Study and Global Meta-analysis

**DOI:** 10.1101/2025.08.18.670785

**Authors:** Kimberley S. Ndlovu, Ricardo R. Pavan, Jacqueline Corry, Ann C. Gregory, Samia Mahamed, Natalia Zotova, Martine Tabala, Pelagie Babakazo, Nicholas T. Funderburg, Marcel Yotebieng, Nichole R. Klatt, Jesse J. Kwiek, Matthew B. Sullivan

## Abstract

Recent studies are revealing that a suboptimal cervicovaginal microbiome (CVMB), including enrichment of anaerobic bacteria associated with multiple female genital disorders, and adverse pregnancy and birth outcomes in pregnant people. Problematically, however, the majority of the available data to date are biased towards highly developed, Global North countries, leaving underrepresented populations like the Democratic Republic of Congo (DRC) poorly characterised. Here, we investigate the CVMB from a cohort of 82 pregnant people living with HIV (PLWH) on antiretroviral therapy (ART) from the DRC. Specifically, we explore the associations between the CVMB via 16S rRNA gene sequencing and maternal peripheral immune factors. Additionally, we compare the CVMB of PLWH-ART from DRC to publicly available CVMB data (5 studies, 1861 samples) in a meta-analysis to elucidate the impact of HIV on the CVMB. Combined, these analyses revealed differences in community structure and predicted function of the microbiota between PLWH-ART and pregnant people without HIV (PWoH). Taxonomically, the CVMB of DRC PLWH-ART were enriched for *Lactobacillus iners-*dominated CVMBs (53%) or a diverse, polymicrobial CVMB, i.e., bacterial vaginosis (BV) (43%). Functional predictions made from these taxa suggested that protein-coupled receptors, amino sugar and nucleotide sugar metabolism, fatty acid metabolism, and polycyclic aromatic hydrocarbon degradation pathways were differentially abundant between communities. Correlation with host plasma immune factors revealed putative links between some CVMB metrics (e.g., alpha diversity and species abundance) that have been linked to adverse pregnancy and birth outcomes.

**Importance:** HIV remains prevalent in sub-Saharan Africa, where it has been linked to adverse birth outcomes. . Suboptimal CVMBs have shown similar links. This pilot study fills critical gaps in understanding how HIV interacts with the pregnant CVMB in populations underrepresented in microbiome research, like the Democratic Republic of Congo. We identified maternal systemic immune factors associated with suboptimal CVMBs that have been linked to poor birth outcomes. In a global meta-analysis, we found significant taxonomic and functional difference in the CVMBs between pregnant people living with and without HIV, revealing potential biomarkers that for increased risks for adverse birth outcomes. These findings provide crucial insights into CVMB features that may influence pregnancy health in pregnant people living with HIV, guiding future research and tailored interventions to support safer pregnancies in the DRC and similar populations.

## Introduction

Human immunodeficiency virus (HIV) is transmitted through sexual contact, blood, needles, or from mother to infant and it attacks immune cells and leads to chronic systemic infection. In July 2024, UNAIDS estimated that women in sub-Saharan Africa account for approximately 62% of HIV infections (1). Several studies have established that pregnant PLWH on ART (PLWH-ART) have a higher risk of adverse pregnancy and birth outcomes, such as low birth weight (2–4) and preterm birth (5–7). This association with adverse birth outcomes also extends to the cervicovaginal microbiome (CVMB), a community of microbes residing in the vagina, which when perturbed towards dysbiosis and bacterial vaginosis (BV) has been associated with preterm birth (8,9) and factors that can lead to preterm birth like premature rupture of membranes (10,11), chorioamnionitis (12,13), and preeclampsia (14).

Currently, the CVMB is characterized using a community state type (CST) framework which can be utilized to assess the compounding impacts of the CVMB and/or HIV infection on pregnancy and birth outcomes. Under this framework, high diversity, non-optimal CVMB, clinically referred to as BV, are classified as CST IV, and *Lactobacillus*-dominated, optimal CVMBs are classified as CSTs I, II, III, and V (15). CST III is the transitional state type that oscillates between being optimal and non-optimal (16). In sub-Saharan Africa, a high percentage of women, particularly those of African ancestry, have CST IV and CST III CVMBs, which are associated with a higher rate of HIV acquisition (17,18) and adverse pregnancy and birth outcomes (19). Therefore, there is a need to elucidate the role of how HIV-associated changes to the CVMB may modulate maternal and infant health outcomes.

The impact of HIV on the CVMB has yielded conflicting results. Some studies report changes in composition, and therefore CST, in the CVMB of PLWH-ART (20,21) while others show no significant differences when compared to people without HIV (PWoH) (22,23). In contrast to these conflicting results, there are some more consistent findings, including that CST IV and HIV-associated CVMBs are linked to increased genital inflammation marked by increased levels of proinflammatory cytokines and chemokines (24,25). In pregnant PLWH-ART such phenotypes are associated with adverse birth and pregnancy outcomes like preterm birth (26–28). Similarly, systemic inflammation PLWH-ART has been well documented (29,30) and is also linked to adverse health outcomes like cardiovascular disease (31,32).

To date, no study has investigated the potential relationship between the CVMB and the soluble systemic immune factors in pregnant PLWH-ART. Here, we present an exploratory study of the CVMB of pregnant PLWH-ART from Kinshasa, Democratic Republic of Congo and to identify CVMB - associated circulating immune factors. Furthermore, we contextualize the results from this with other CVMB datasets that include pregnant people in a meta-analysis, providing a more global perspective on the interactions between the CVMB and PLWH-ART. Finally, we discuss how these findings relate to pregnancy and birth outcomes in PLWH-ART.

## Results and discussion

### CQI-PMTCT study population, sampling strategy, data generation and meta-analysis

In this pilot study, we sought to generate 16S rRNA gene sequencing data to investigate the CVMB of PLWH-ART in the CQI-PMTCT cohort and its association to the systemic immune system. To that end, we collected cervicovaginal swabs and peripheral blood from a subset of 82 pregnant PLWH-ART living in Kinshasa, DRC to generate 16S rRNA gene data and plasma immune factor levels, respectively (see **Methods, Fig. 1A**). Participant characteristics, stratified by CST, including participant age, gestational age, ART regimen, gravidity, socioeconomic status, and adverse birth outcomes are summarized in **Table 1**. The CQI-PMTCT cohort included only PLWH-ART participants, and since pregnant PWoH groups were lacking for comparison, the conclusions we can make from these analyses are limited. Hence, we also conducted a meta-analysis where we integrated CVMB data from the CQI-PMTCT cohort with other published and publicly available pregnant CVMB datasets to investigate the differences in the CVMB between PLWH-ART and PWoH (**Fig. 1B**).

**Fig 1.**
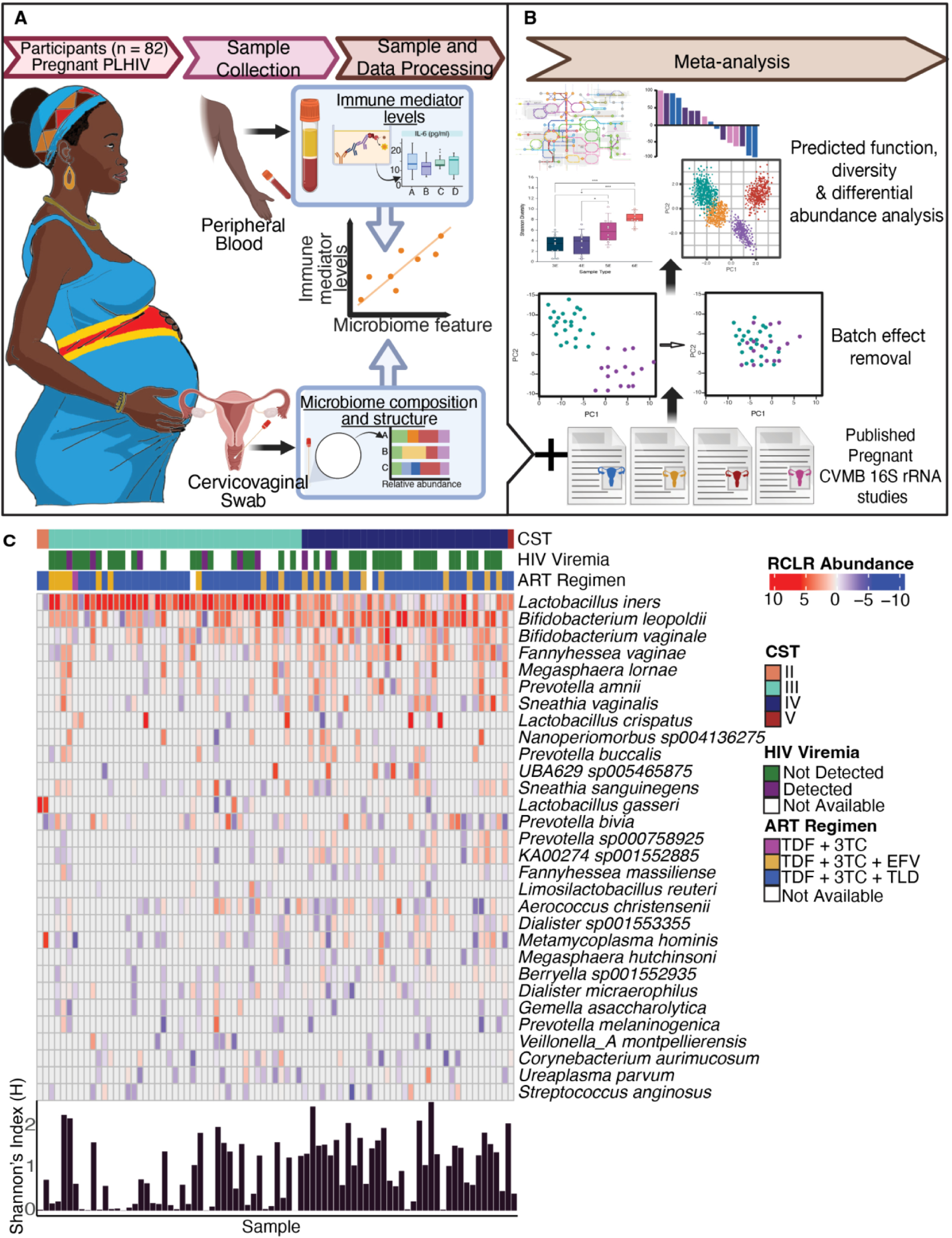
Study schematic and CVMB composition of the CQI-PMTCT cohort. **A)** Study schematic of the CQI-PMTCT study. Peripheral blood and cervicovaginal swabs were collected from PLWH-ART in the Democratic Republic of Congo and processed in Columbus, OH. **B)** CVMB data from the CQI-PMTCT cohort were combined with other publicly available pregnant CVMB data in a meta-analysis **C)** Heatmap of RCLR abundances of top 30 most abundant microbial taxa from 81 women. CSTs were determined using VALENCIA. HIV viremia, ART regimen, and per sample Shannon Index values are shown.

**Table 1:**
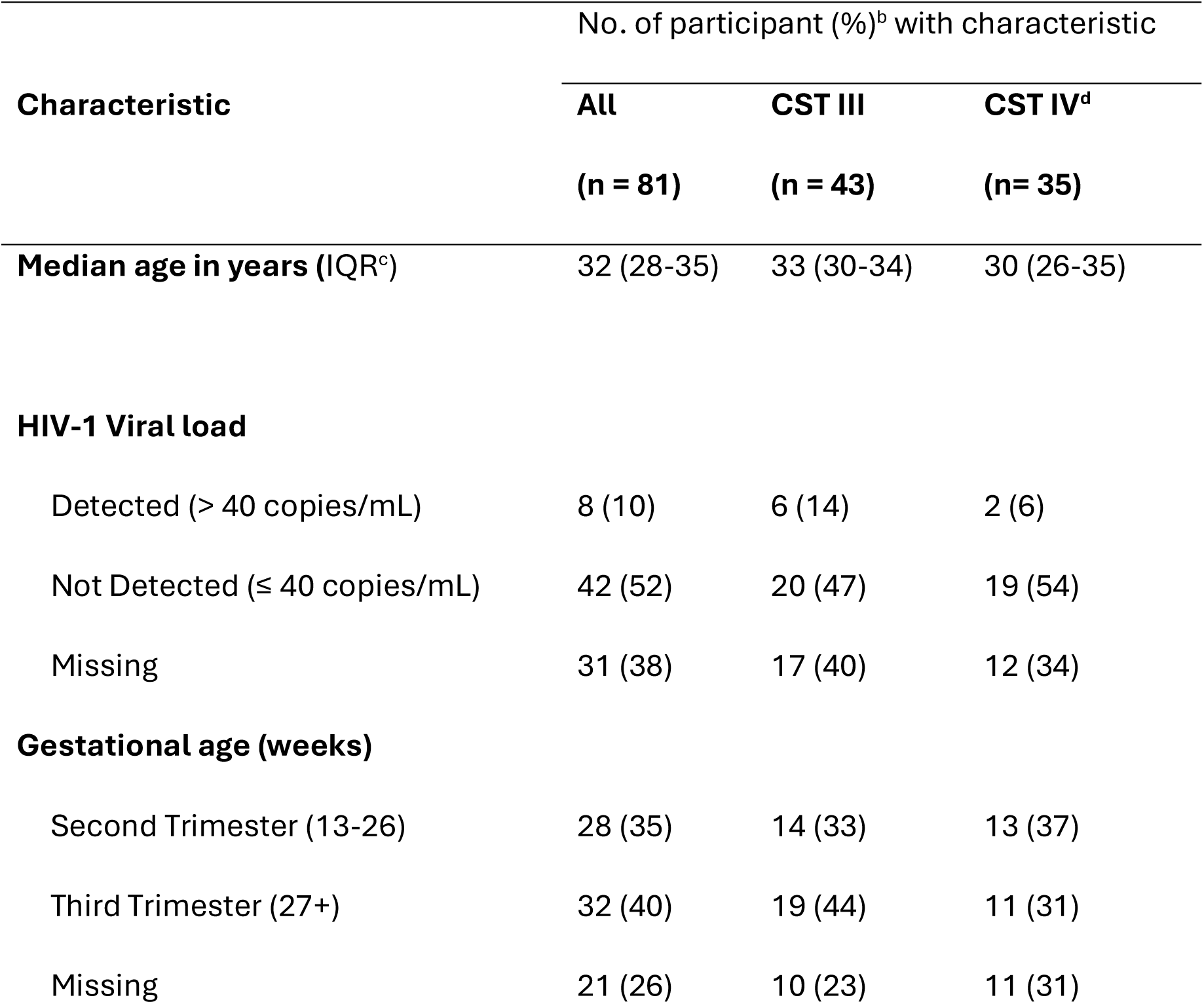
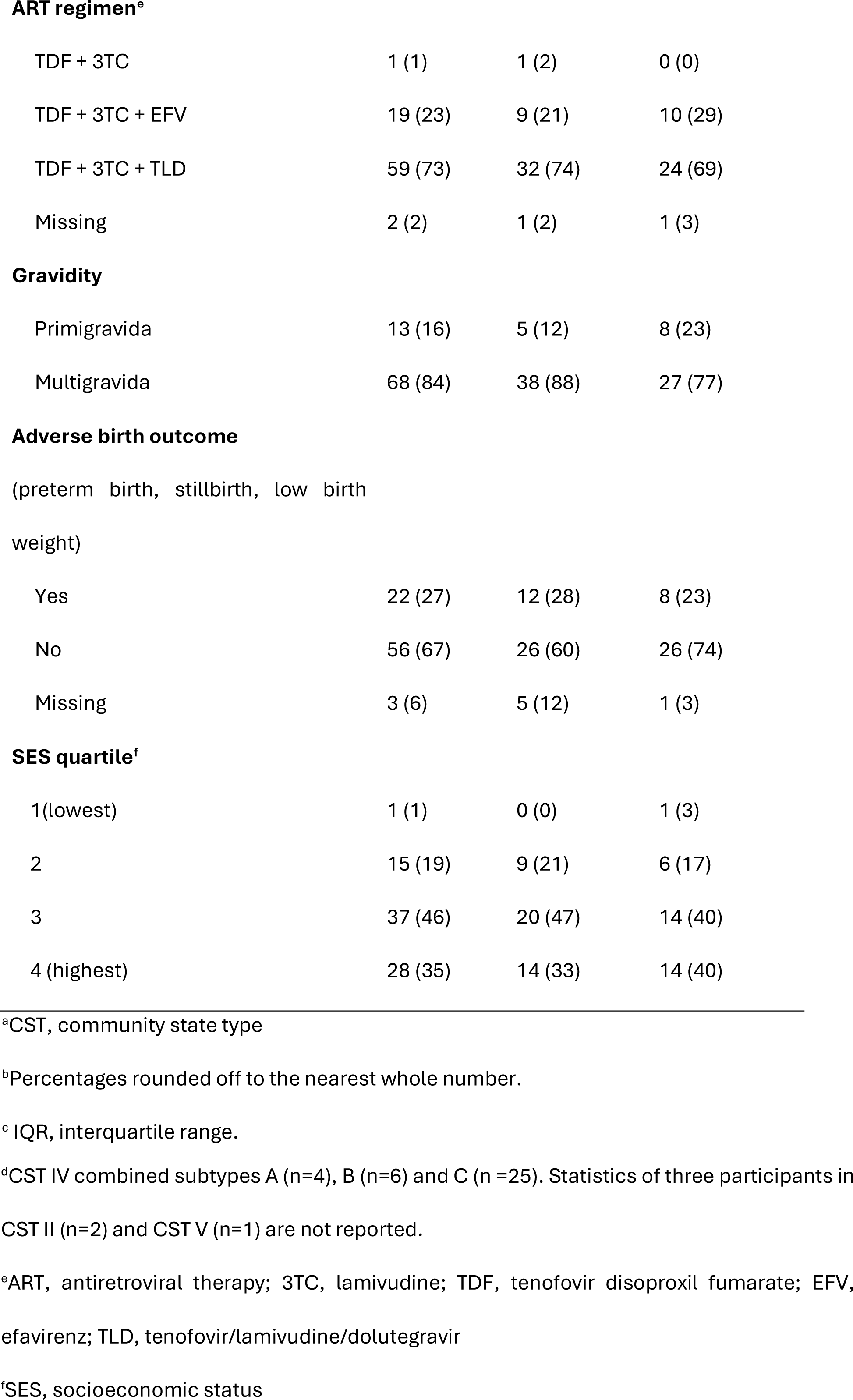
Participant demographics at enrolment stratified by community state type^a^.

| Characteristic | No. of participant (%)<sup>b</sup> with characteristic |  |  |
| --- | --- | --- | --- |
|  | All | CST III | CST IV<sup>d</sup> |
|  | (n = 81) | (n = 43) | (n = 35) |
| **Median age in years (IQR<sup>c</sup>)** | 32 (28-35) | 33 (30-34) | 30 (26-35) |
| **HIV-1 Viral load** |  |  |  |
| Detected (> 40 copies/mL) | 8 (10) | 6 (14) | 2 (6) |
| Not Detected (≤ 40 copies/mL) | 42 (52) | 20 (47) | 19 (54) |
| Missing | 31 (38) | 17 (40) | 12 (34) |
| **Gestational age (weeks)** |  |  |  |
| Second Trimester (13-26) | 28 (35) | 14 (33) | 13 (37) |
| Third Trimester (27+) | 32 (40) | 19 (44) | 11 (31) |
| Missing | 21 (26) | 10 (23) | 11 (31) |
| **ART regimen<sup>e</sup>** |  |  |  |
| TDF + 3TC | 1 (1) | 1 (2) | 0 (0) |
| TDF + 3TC + EFV | 19 (23) | 9 (21) | 10 (29) |
| TDF + 3TC + TLD | 59 (73) | 32 (74) | 24 (69) |
| Missing | 2 (2) | 1 (2) | 1 (3) |
| **Gravidity** |  |  |  |
| Primigravida | 13 (16) | 5 (12) | 8 (23) |
| Multigravida | 68 (84) | 38 (88) | 27 (77) |
| Yes | 22 (27) | 12 (28) | 8 (23) |
| No | 56 (67) | 26 (60) | 26 (74) |
| Missing | 3 (6) | 5 (12) | 1 (3) |
| **SES quartile<sup>f</sup>** |  |  |  |
| 1(lowest) | 1 (1) | 0 (0) | 1 (3) |
| 2 | 15 (19) | 9 (21) | 6 (17) |
| 3 | 37 (46) | 20 (47) | 14 (40) |
| 4 (highest) | 28 (35) | 14 (33) | 14 (40) |

<sup>a</sup>CST, community state type
<sup>b</sup>Percentages rounded off to the nearest whole number.
<sup>c</sup> IQR, interquartile range.
<sup>d</sup>CST IV combined subtypes A (n=4), B (n=6) and C (n =25). Statistics of three participants in CST II (n=2) and CST V (n=1) are not reported.
<sup>e</sup>ART, antiretroviral therapy; 3TC, lamivudine; TDF, tenofovir disoproxil fumarate; EFV, efavirenz; TLD, tenofovir/lamivudine/dolutegravir
<sup>f</sup>SES, socioeconomic status

### In the CQI-PMTCT cohort, CST III and CST IV types are predominant in the CVMB of pregnant PLWH-ART in the DRC

To date, no study has investigated the CVMB of people in the DRC. We generated 16S rRNA gene sequencing data and survey the CVMB of pregnant PLWH-ART. Focusing on the top 30 most abundant species for CVMB analyses revealed 53% of the participants (n = 43) were dominated by *Lactobacillus iners,* while another 43% of the participants (n=35) were dominated by a consortia of polymicrobial anaerobic bacteria (i.e., *Bifidobacterium leopoldii , Bifidobacterium vaginae, Fannyhesea vaginae, Prevotella spp.* and *UBA629 sp0054587)* (**Fig. 1C**). These findings classify the CVMBs of PLWH-ART to CST III (transitional CST) or CST IV (the non-optimal CST), respectively.

Evaluating diversity across these datasets revealed the following. First, within sample diversity showed significant differences between identified CSTs (p = 0.0023, Kruskal-Wallis) (**Fig. 2A**). This agrees with the paradigm that CST IV CVMB are more diverse compared to *Lactobacillus-*dominated CSTs (15). Second, alpha diversity demonstrated that samples cluster by CST (ANOSIM, R = 0.206, p = 0.001, **Fig. 2B**), with *L. iners* driving the highest variation in the CST III cluster and *B. vaginale, B. leopoldii and F vaginae,* are driving the variation in the CST IV cluster.

**Fig 2.**
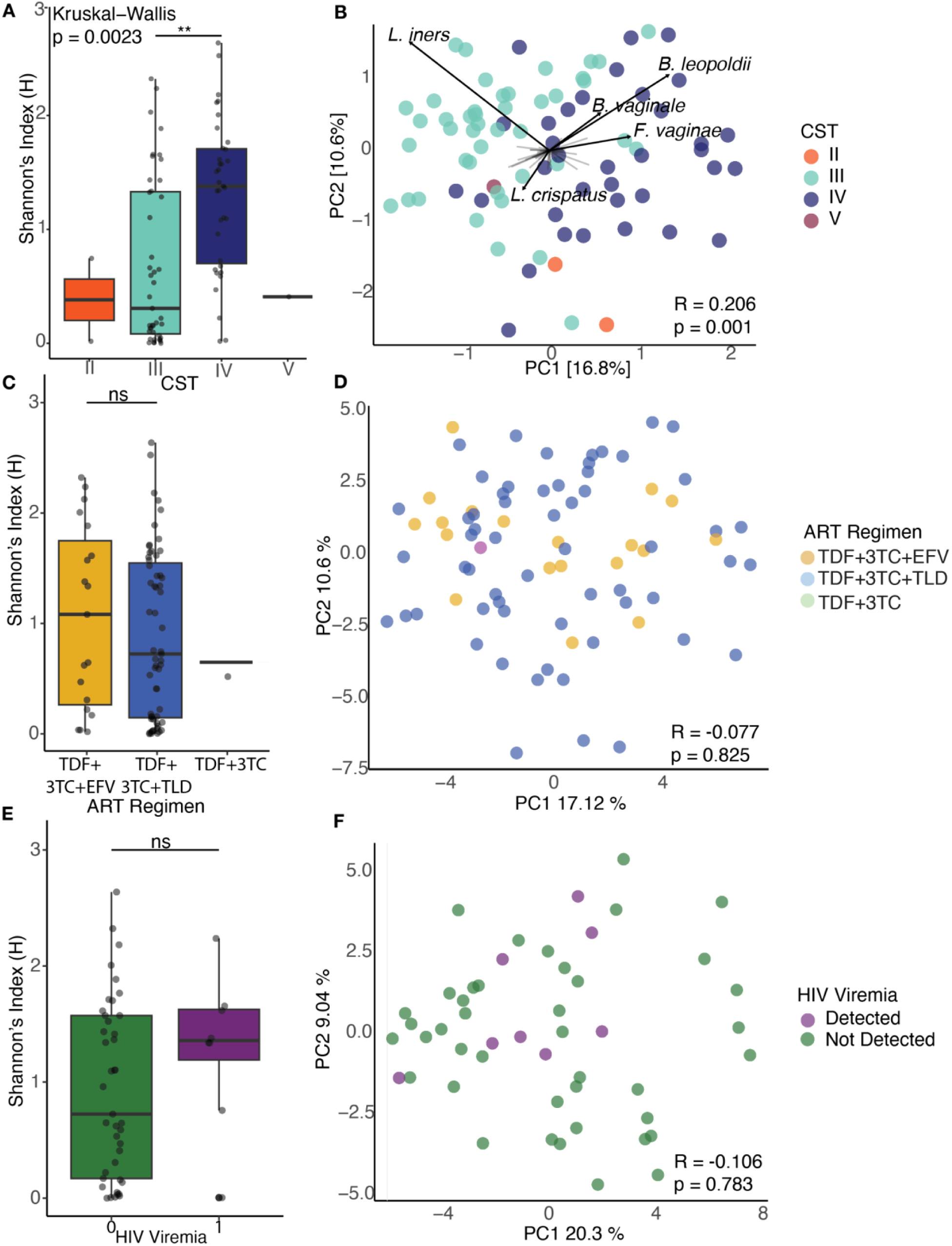
Ecological statistics of the CVMB in the CQI-PMTCT cohort. **A)**. Shannon entropy diversity stratified by CSTs (Kruskal-Wallis p =0.00234). **B)** PCA biplot of Robust Aitchison distances coloured by CSTs (ANOSIM, R = 0.206,p = 0.001). The loadings of the top 5 most abundant species across all samples are labelled to primary highlight drivers of variation amongst CST clusters with black arrows to indicate the magnitude and directionality. **C)** Shannon indices aggregated by ART regimen (Kruskal-Wallis p=0.695). **D)** PCA of Robust Aitchison distances colored by ART regimen for 79 participants (ANOSIM R =-0.077, p = 0.825). **E)** Aggregated Shannon entropy values of HIV viremia for 50 participants (Wilcox test p = 0.2853). **F)** PCA of robust Aitchison distances colored by detected viral load (ANOSIM R= -0.106, p = 0.783). Within each boxplot, the horizontal lines denote median values and the boxes extend from the 25^th^ to the 75^th^ percentile. The vertical lines in each box plot show the values with 1.5 IQR. **p<0.01, ns>0.05

These results are consistent with other studies that show sub-Saharan African women are more likely to harbor a high risk, non-optimal CST IV CVMB (17,33,34). Additionally, other studies with similar cohorts in geography and size have shown that pregnant PWH have CVMBs largely dominated by *Gardnerella spp.* and *L. iners* (21,28). Incidentally, *L. iners* is the common *Lactobacilli* in CVMBs of sub-Saharan African people and people across the globe (15,33), however it is not the most optimal *Lactobacillus* compared to *L. crispatus* or *L. jensenii*.

Despite its high prevalence in the global CVMB, the role of *L. iners* in vaginal health and in pregnancy and birth outcomes remains ambiguous. For example, in pregnant PLWH-ART, *L. iners*-dominated CVMBs were associated with risk of preterm birth (21,28), however, in pregnant PWoH, *L. iners* was associated with lower risk of preterm birth (35,36). These studies underscore the complexity of *L. iners* role in vaginal health and pregnancy outcomes. *L. iners* can exist with other ‘non-optimal’ CST IV bacteria, is less stable in the vaginal microbiome and so can easily allow the transition to non-*Lactobacillus* dominated, CST IV communities (i.e. BV) (37). Additionally, *L. iners* has been shown to have pathogenic potential; for example, it produces the less protective L-lactic acid isomer (38–40), and has highly expressed homologs of cholesterol-dependent lysins, pore forming exotoxins, which have been found in other pathogenic bacteria and not in other *Lactobacilli* (41–43). However, its presence even in healthy, asymptomatic people, along with its existence as a variety of strains, demonstrates that it is also a vaginal symbiont and is able to persist in the CVMB despite other more beneficial or protective vaginal *Lactobacilli* (44,45). Further studies are needed to clarify the enigmatic roles of *L. iners* in vaginal health and pregnancy outcomes using strain resolved metagenomics.

The other two most abundant bacteria in the CQI-PMTCT cohort, *B. leopoldii* and *B. vaginale* (both formerly *G. vaginalis* until a recent amendment (46)), have been shown to be in high abundance in sub-Saharan populations (21,28), even in healthy, asymptomatic CVMBs (47–49). These CST IV bacteria have been associated with adverse pregnancy and birth outcomes like preterm birth (8,50) and fetal death (51–53). Overall, our results are in accordance with other studies showing the CVMBs of sub-Saharan African PLWH-ART have non-*Lactobacillus* dominated (CSTs IV) or *L.iners*-dominated (CST III) CVMBs, and may contribute to higher rates of HIV and sexually transmitted infections (54,55) as well as preterm birth (20,21).

### CVMB structure is not associated with HIV viremia and ART regimen in the CQI-PMTCT cohort

Next, we sought to investigate clinical characteristics that might influence the CVMB in the CQI-PMTCT. We focused particularly on type of ART regimen and HIV viremia. Currently, there are inconclusive results on whether ART impacts the CVMB (56–58). The same is true for HIV viremia, particularly in chronic HIV infection. (21,22,59). Therefore, we asked whether the type of ART regimen or HIV viremia in PLWH-ART had influence in the diversity and structure of the CVMB. We hypothesized that type of ART will impact the CVMB and that participants with detectable HIV viremia (n = 8, **Table 1**) will have more diverse CVMB, with communities that were distinct from those with un-detectable HIV viremia (n =42, **Table 1**).

Contrary to our hypotheses, we detected no significant differences in alpha diversity at the species level between ART regimen (Kruskal Wallis test p = 0.695, **Fig. 2C**) and no differences in CVMB structure between types of ART regimen (ANOSIM R= -0.077, p = 0.825, **Fig. 2D**). We also did not see differences in alpha diversity between detected and undetected HIV viremia (Wilcoxon Test, p =0.285, **Fig. 2E**) and in beta diversity (ANOSIM R= -0.106, p = 0.783, **Fig. 2F**).

Studies with similar cohorts in size and geographic location have shown that ART use did not impact the CVMB of Ugandan PLWH-ART (56). However, CVMBs have been associated with varying concentrations of ART in the female genital tract and plasma meaning there is a possibility that long term ART use could have some influence in the CVMB (60). Currently, it is known that topical ARTs, like PreP and tenofovir, are metabolized by CST IV cervicovaginal bacteria thereby reducing their efficacy (61–63) but these ARTs do not seem to alter the CVMB composition and structure (56–58). While further research might be necessary to investigate the link between ART use and CVMB composition, our results contribute to the growing knowledge of the impact, or lack thereof, between ART use and the CVMB (64).

Studies have shown before that there are no differences in vaginal communities due to HIV viremia or status (22,59). Conversely, other 16S rRNA gene studies have shown that HIV-associated CVMBs have a lower abundance of *Lactobacillus* species but there was no conclusion on how CVMB structure (i.e. beta diversity) is different (55,65,66). Curiously, there is inconclusive evidence on how HIV influences CVMB composition. On one hand, cervicovaginal bacteria have been shown to interact with the HIV virus or host cells to increase risk of contracting HIV. For example, CST IV bacteria like *G. vaginalis* induce a local inflammation response, reduce cervicovaginal epithelial cell barrier and integrity hence allowing bacterial and viral pathogens, like HIV, to infect the host (17,67,68). A recent study also showed that bacterial lectins specific to O-glycans, bind to the HIV-1 virion and envelope glycoproteins to increase HIV infectivity and resistance to antibodies (69). However, the mechanisms by which HIV directly impact bacterial physiology and metabolism remain undefined and unexplored. It is possible that the HIV viral load in the vagina is sufficiently suppressed for there to be a detectable interaction between the virus and vaginal bacteria (70,71), however studies on this subject are highly limited. The ambiguity of the impact of chronic and systemic viral sexually transmitted infections like HIV on the CVMB remains to be fully explored however, based on our results for the CQI-PMTCT cohort, we can tentatively conclude and recapitulate the findings of studies that state HIV viremia does not influence the CVMB (22,23,59). The high percentage of people with undetectable HIV viremia is a testament to the effectiveness of ART use in this cohort.

### Meta-analysis reveals that CVMB are distinct between PLWH-ART and PWoH globally

The CQI-PMTCT study discussed above has limitations including lack of a pregnant PWoH group for comparison. Hence, to better understand the role of HIV on the CVMB, we integrated this study with five other publicly available pregnant CVMB datasets. To reduce batch effects, data from studies were selected to be as similar to the CQI-PMTCT CVMB data as possible (see **Methods** and **Fig. S1**).

After adjusting for study specific batch effects using PLSDA (see **Methods**), we found that within sample (alpha) species diversity was significantly higher in samples from PLWH-ART compared to PWoH (**Fig. 3A**) Additionally, CVMB community structure was significantly different between pregnant PLWH-ART and pregnant PWoH CVMBs (ANOSIM R= 0.518, p =0.001, **Fig. 3B**). When stratified by CST, within samples species diversity was highest in CST IV samples, which is suboptimal in the CVMB (**Fig. 3C**) and CVMB community structure was different by CSTs (ANOSIM R= 0.399, p =0.001, **Fig. 3D**). All this indicates that there are different bacterial communities between the PLWH-ART and PWoH groups.

**Fig 3.**
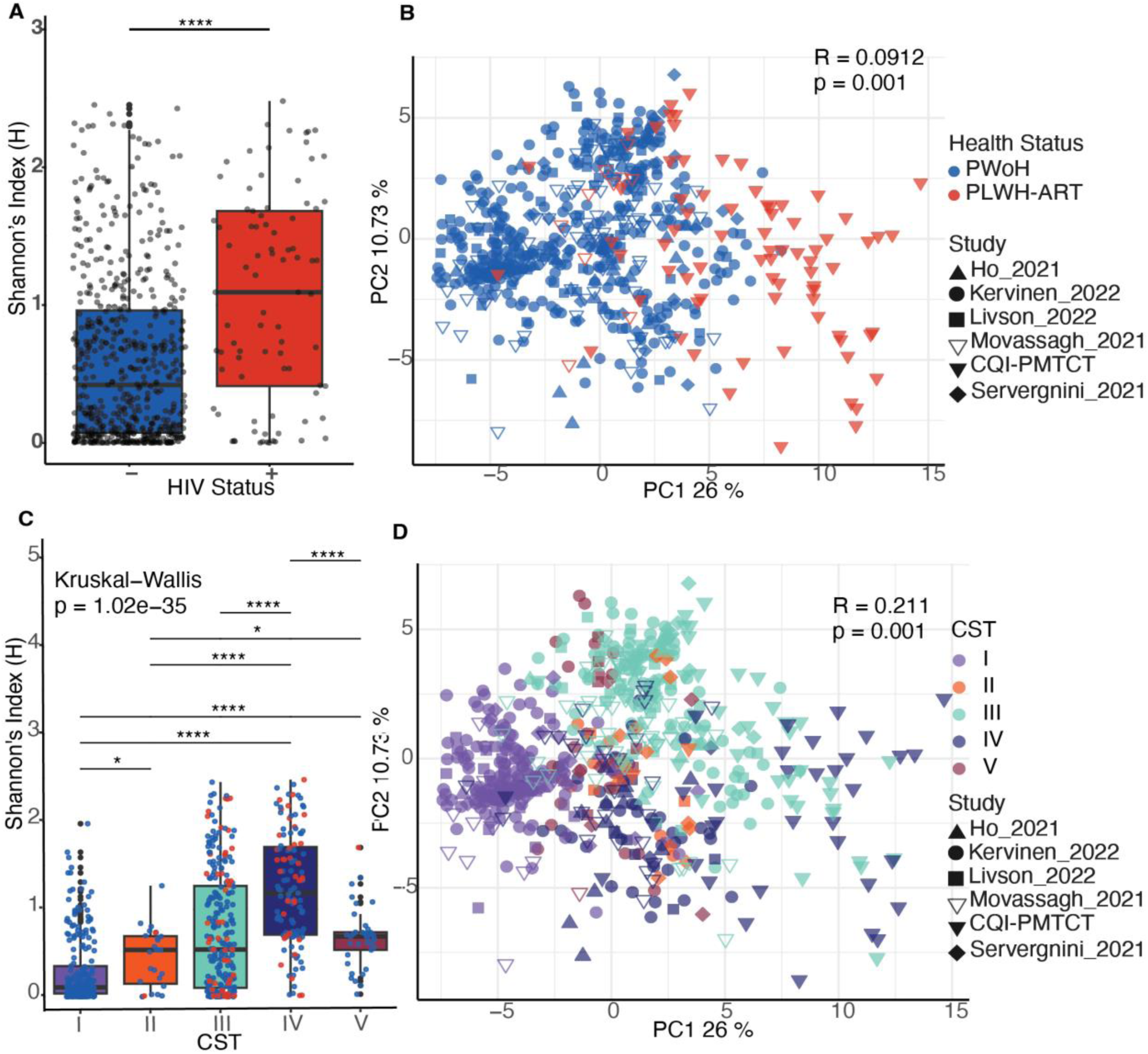
Ecological statistics of the integrated datasets. **A)** Shannon Entropy values between CVMBs of PWoH and PLWH-ART separated by study. **B)** PCA of Aitchison Distances showing different bacterial communities by health status (ANOSIM R= 0.518, p= 0.001). **C)** Shannon Entropy values stratified by CST (Kruskal-Wallis p = 1.023e-35). **D)** PCA of Aitchison Distances showing different bacterial communities by CST (ANOSIM R=0.399, p= 0.001). *p<0.05, ****p<10^-4^

To further investigate this claim, we used differential abundance analyses (see **Methods**) to identify bacterial species that were differentially enriched between the two groups. We found that 21 cervicovaginal bacterial species from 14 families were differentially abundant between the PLWH-ART and PWoH groups (**Fig. 4**). *Lactobacilli spp.,* namely beneficial bacteria such as *L. crispatus, L. gasseri, L. hominis* and *L. jensenii* and *Limosilactobacilus reuteri* were more abundant in the CVMB of PWoH. In contrast, the PLWH-ART group had high abundance of CST IV, non-optimal, BV-associated bacteria compared to the PWoH group. For example, species in the *Bacteroidaceae* family, particularly *P. amnni* and *P.timonensis spp.* were more abundant in the PLWH-ART group. Other CST IV associated bacteria that were more abundant in the PLWH-ART group were *B. leopoldii, M. lornae, KA00274 sp001552885* (AKA *Amygdalobacter nucleatus*), *F.magna, F. vaginae, M. hominis* and *C. tuberculostearicum* (**Fig. 4**).

**Fig 4.**
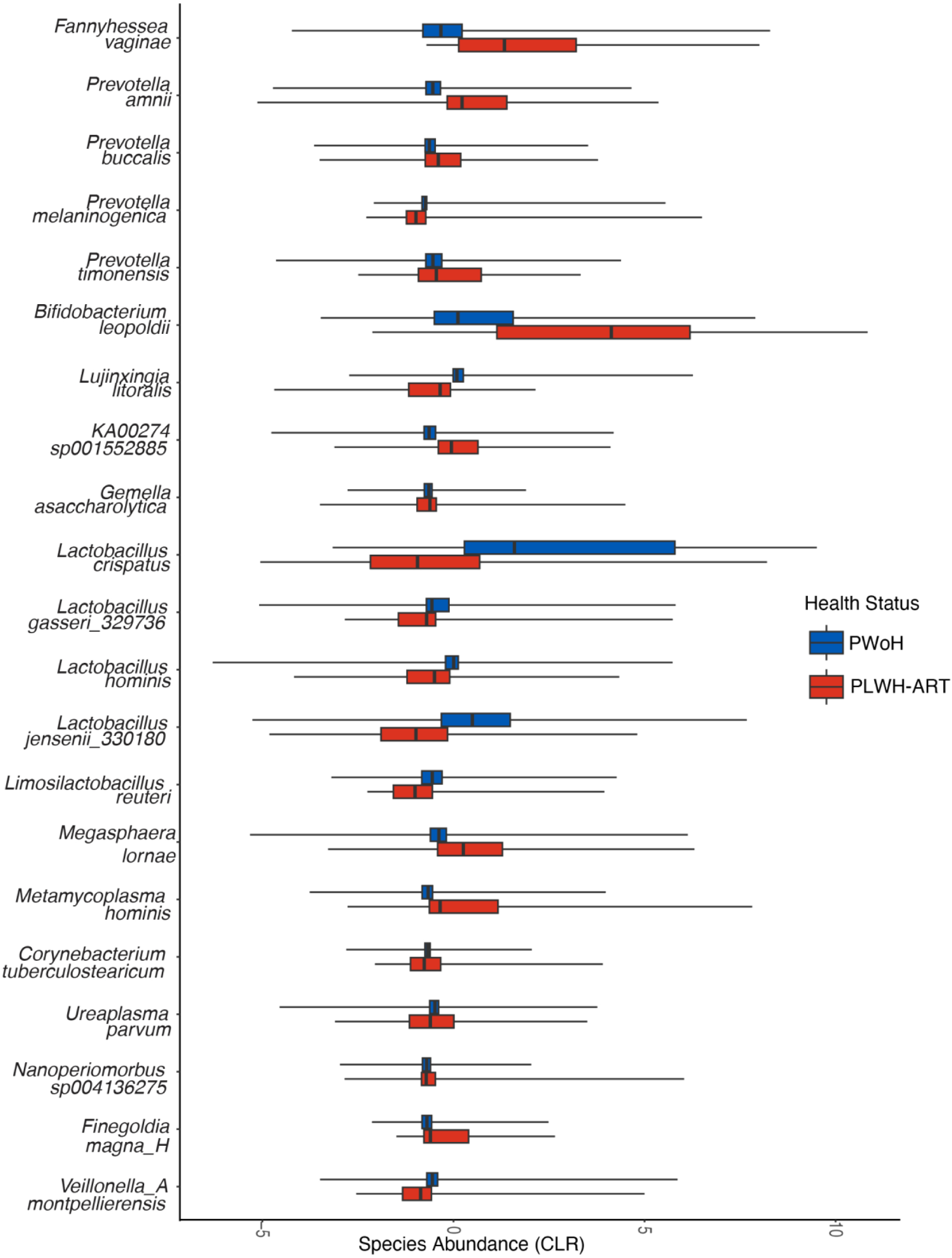
Boxplots showing differentially expressed species in pregnant PLWH-ART and pregnant PWoH as determined using three differential abundance analysis methods (see **Methods)**. Within each box, the vertical lines denote median values, and the boxes extend from the 25^th^ to the 75^th^ percentile. The horizontal lines in each box plot show the values with 1.5 IQR.

Other studies have found *Lactobacillus spp.* to be more dominant in PLWH not on ART compared to those on ART, regardless of pregnancy state (55,65,66). This is corroborated by the CST IV, non-optimal bacteria being more differentially abundant in PLWH-ART group compared to the PWoH. It is possible that this differential abundance in (non)- *Lactobacillus-*dominated CVMBs between PLWH-ART and PWoH existed prior to and during HIV acquisition, as such differences are associated with risk of HIV acquisition (72). For that reason, these communities may have remained stable in their respective states since. Indeed, one study determined that in 79.7% of CVMBs from two longitudinal cohorts (16,73), CSTs were predominantly mono-stable and rarely transitioned to completely different CSTs, which could explain the persistence of these bacteria in the CVMB even after HIV acquisition (37). It is also possible that the influence of maternal age, racioethnicity and geography (**Fig. S2**) distorts the impact of HIV status on the CVMB in this meta-analysis as seen before (47,48,74,75). Therefore, we tentatively conclude that HIV infection is associated with changes in the CVMB of pregnant people as seen in a recent study with similar populations (76). However, future research will have to explore the causative associations of HIV on the pregnant CVMB and how this alters pregnancy outcomes.

### Predicted metabolic pathways were differentially abundant among CSTs in the CQI-PMTCT cohort

The CST framework classifies the CVMB based on taxonomic composition (15). However, even when microbial communities differ in their taxonomic composition, they might have functional redundancy (77) or conversely, diverse functional capabilities in the same species (78). Hence, functional prediction of the CVMB may yield more specific insights into potential phenotypic differences between CST, and thus host responses. Here, we predicted the function of the CVMB, stratified by CST, using PICRUSt2 based on the KEGG (Kyoto Encyclopedia of Genes and Genomes) database (79,80). We predicted 6466 KEGG Orthologs (KOs), which were assigned to 162 KEGG pathways. Of note, these are predicted functions and not direct readouts by the bacterial species.

Using multiple differential abundance analyses (see **Methods**), we identified four (4) KEGG pathways that were differentially abundant between CSTs (**Fig. 5**), namely G protein-coupled receptors (GPCRs, KO04030), amino sugar and nucleotide sugar metabolism (KO00520), fatty acid metabolism (KO01212) and polycyclic aromatic hydrocarbon degradation (KO00624). Fatty acid metabolism and polycyclic aromatic hydrocarbon degradation had higher relative abundance in CST IV compared to CST III, with the highest abundance of both in CST II. G protein-coupled receptors and amino sugar and nucleotide sugar metabolism were more abundant in CST III, followed by CST IV and lastly CST II (**Fig. 5**).

**Fig 5.**
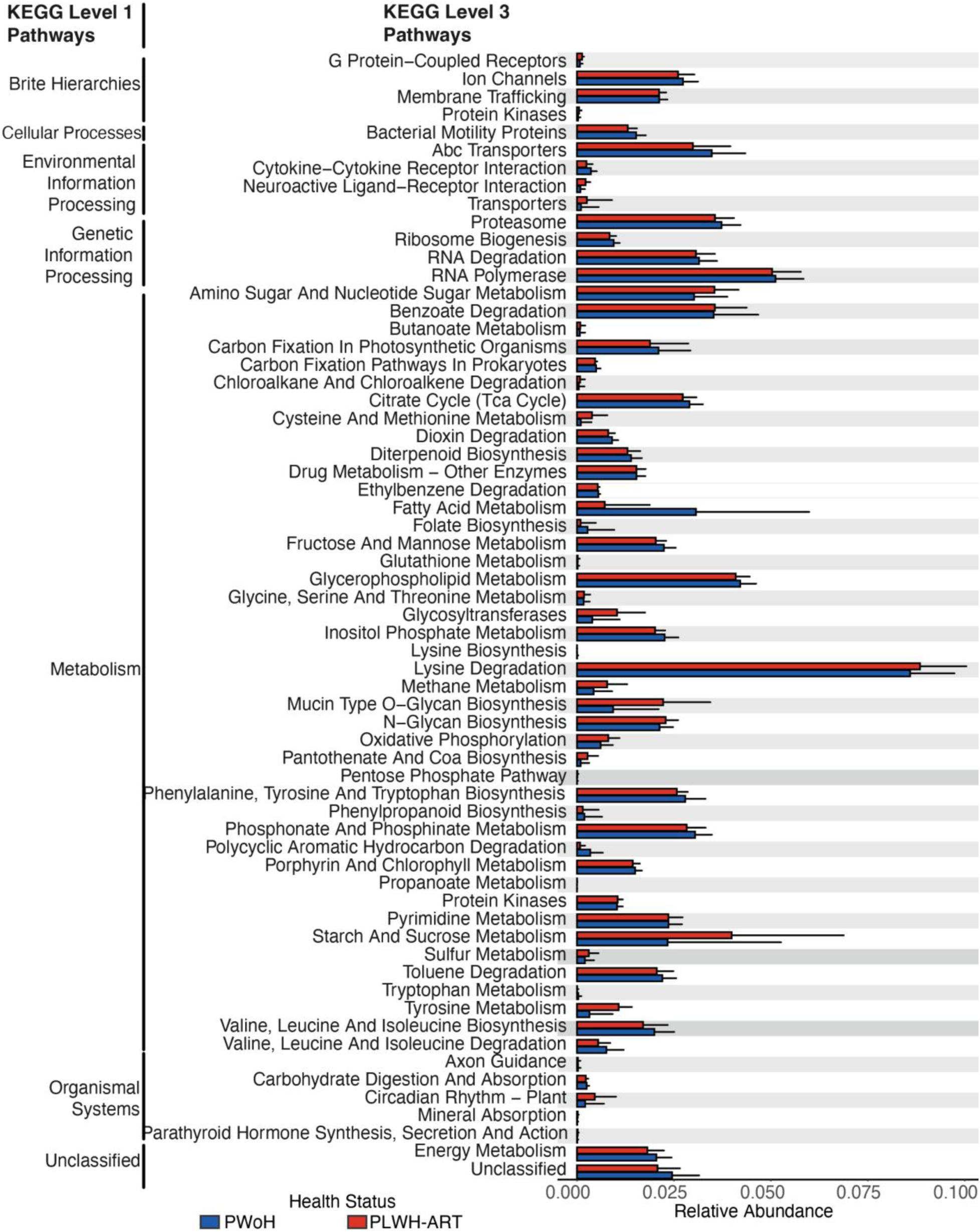
Predicted KEGG pathways differentially abundant between CSTs in the CQI-PMTCT cohort as determined using three differential abundance analysis methods (see **Methods)**.

Because there were only two samples that were assigned CST II, we will focus our discussion on CST III and IV only. A combination of predicted CVMB genotypes suggests a plausible mechanism for disruption of host homeostasis. Short-chain fatty acid (SFCA) metabolism was more abundant in CST IV compared to CST III CVMBs, indicating greater presence of bacteria capable of producing SFCAs. CST IV bacteria play a large role in SCFA production, and the increased presence of SFCAs like propionate, acetate and butyrate have been associated with cervicovaginal dysbiosis (81). Furthermore, the production of SCFA leads to decreased *Lactobacillus* abundance and increased pH in the vagina (81–83). Based on this, we can posit that there were high concentrations of cervicovaginal SFCAs in this cohort. GPCRs were slightly, yet still significantly, more abundant in CST III compared to CST IV CVMSs. GPCRs mediate the immunoregulatory effects of SFCAs (82) and specifically, GPCRs 41, 43 and 109A, can inhibit histone deacetylases (HDACs) which subsequently induce a cellular and humoral immune response (82,84). Simultaneously increased SFCAs and decreased GPCRs may induce a local pro-inflammatory response in this cohort. Given that inflammation in the cervicovaginal tract is associated with adverse birth outcomes, like preterm birth in pregnant PLWH-ART (21), which may be linked to increases in SFCA production and decreases in the immunoregulators in the CVMB.

Amino sugar and nucleotide sugar metabolism was another pathway that might be implicated in birth outcomes in pregnant PLWH-ART. For bacteria, these are core intermediates involved in peptidoglycan and lipopolysaccharide, among other macromolecule, biosynthesis, energy metabolism, and other processes. In this cohort, this pathway was more abundant in CST III compared to CST IV CVMBs. One study found that amino sugar and nucleotide metabolism was negatively associated with mammalian target of rapamycin (mTOR), a kinase that regulates cell growth and proliferation (85). In the same study, mTOR was negatively correlated with *L. iners* and positively correlated with polymicrobial, CST IV bacteria (85). While some of this association may be due to major structural features of the CVMB, i.e., the presence of a lipopolysaccharide layer of some members but not others, these metabolic products from the CVMB of pregnant PLWH-ART may interact with the mTOR pathway, activating it (by CST III bacteria) or inhibiting it (by CST IV bacteria). This interaction could maintain or disrupt cervicovaginal epithelial integrity, potentially determining whether birth outcomes are adverse or normal through a proinflammatory or anti-inflammatory response and either increasing or decreasing the risk of STIs.

Lastly, polycyclic aromatic hydrocarbon (PAH) degradation was also differentially more abundant in CST IV than in CST III CVMBs. PAHs, which are ubiquitous in the environment can impact the CVMB (9) which can lead to adverse birth outcomes like intrauterine growth restrictions and preterm birth (86,87). The polymicrobial state type, CST IV, tends to have varying abundances of putative PAH-degrading bacterial genera including *BVAB2* and *Megasphaera, Sphingomonas*, *Acinetobacter*, *Micrococcus*, *Pseudomonas*, and *Ralstonia,* and two of the genera, *Megasphaera* and *Pseudomonas* are present in the CQI-PMTCT cohort (88–90). Therefore, somewhat unexpectedly, a more diverse, polymicrobial CST IV CVMB that may be better able to degrade PAHs might offer some protection against their harmful effects. Perhaps, because the Sub-Saharan African region is exposed to a substantial amount of PAHs (91,92), a CST IV CVMB might be advantageous and help explain its prevalence in Sub Saharan African populations (17,48,93). However, a multidisciplinary research study, across life and social sciences, would need to be conducted to investigate this claim (94,95), which is an important next step. Overall, these findings provide the direction for further studies to clarify these differences with more granularity and the role of bacterial metabolic pathways on reproductive and birth outcomes in pregnant PLWH-ART in vulnerable and underrepresented populations. Additionally, future multi-omics studies should be conducted to progress the CST framework from taxonomy-only based classification to microbial function and taxonomy-based classification.

### Meta-analysis reveals predicted metabolic pathways were differentially abundant between PLWH-ART and PWoH

We used PICRUSt2 to predict 4,534 KOs which were assigned to 130 KEGG pathways. Of those, 62 KEGG pathways were differentially abundant between pregnant PWoH and PLWH-ART CVMBs (**Fig. 6**).

**Fig 6.**
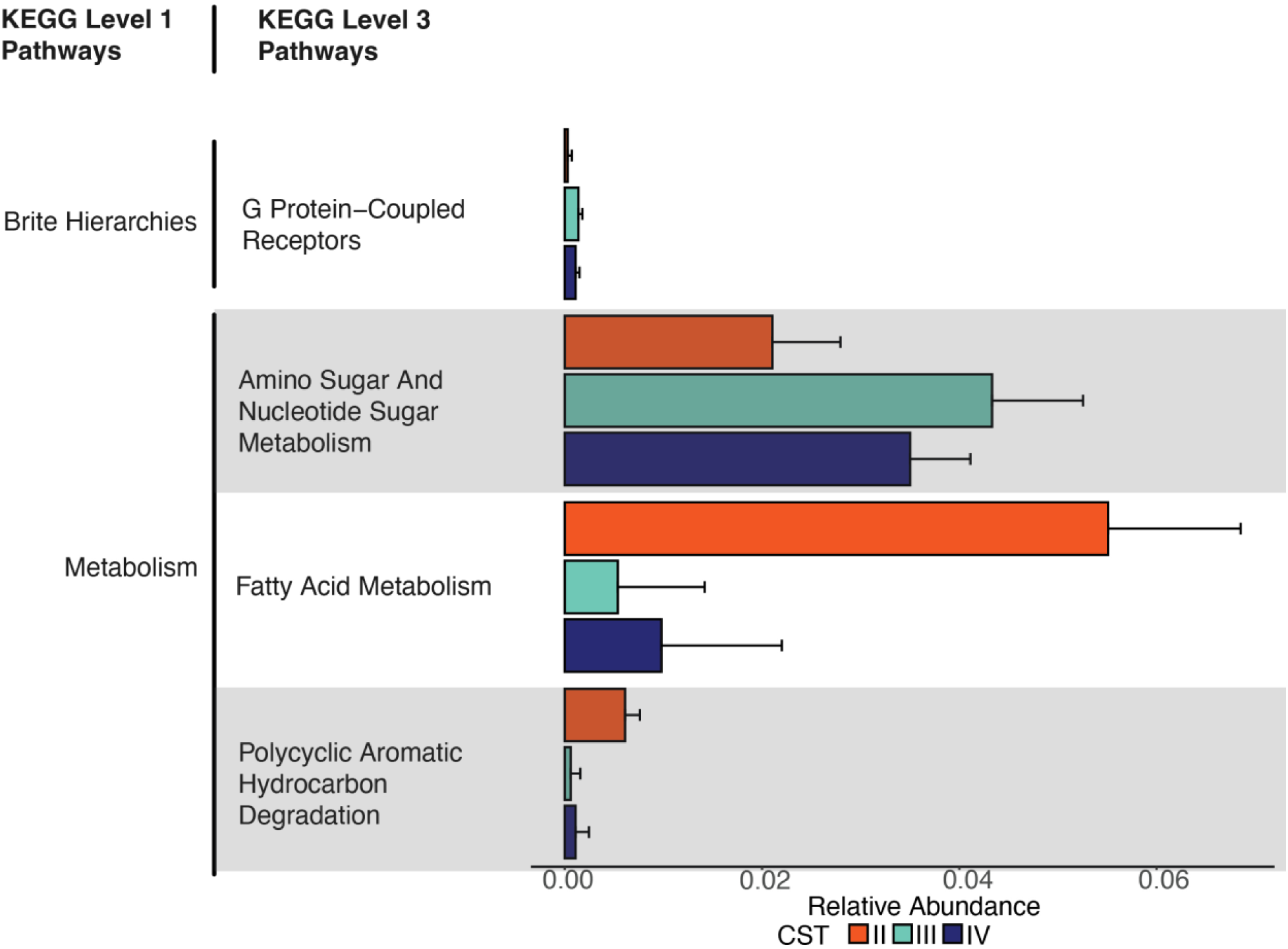
KEGG pathways differentially abundant between pregnant PLWH-ART and pregnant PWoH as determined using three differential abundance analysis methods (see **Methods)**.

Most of these pathways were primarily involved in metabolism. This could indicate that HIV-associated changes to the pregnant CVMB may alter microbial function, which could influence pregnancy and birth outcomes and susceptibility to other cervicovaginal infections in PLWH-ART. For this study, we will focus on only a few pathways that might have relevance with HIV, and pregnancy and birth outcomes. Lysine degradation (KO00310) was differentially more abundant in the PLWH-ART group compared to the PWoH group. Products of lysine degradation — cadaverine and pipecolate — were increased in cervicovaginal fluids of women with BV (96). BV, indicative of CST IV, has been associated with adverse birth outcomes in PLWH-ART (20,21,28) and increased risk of HIV acquisition (17). Cysteine and methionine metabolism (KO00270) was differentially more abundant in the PLWH-ART group compared to the PWoH group, and the opposite is true for folate biosynthesis (KO00790). Levels of cysteine in maternal plasma or cervicovaginal fluid have been associated with adverse pregnancy and birth outcomes (97–99). These findings provide direction for further research to investigate how and if HIV-associated changes to the CVMB function impact pregnancy and birth outcomes. One pathway, mucin O-glycan synthesis (KO00512) was differentially more abundant in PLWH-ART and this has been shown to increase the infectivity of sexually transmitted viral pathogens like HIV (69). Future research should investigate how this pathway is involved in pregnancy and birth outcomes in PLWH-ART.

### CVMB characteristics are correlated with some plasma immune factors in the CQI-PMTCT cohort

Next, we asked if there was an association between plasma immune factors and the CVMB in the CQI-PMTCT cohort. The correlation and causative associations between CVMB and cervicovaginal cytokines and chemokines have been well studied (17,24,67,100). Few studies have identified the link between the systemic immune system and the CVMB in PLWH-ART. To this end, we used linear models to identify the CVMB metrics (Shannon diversity, CST and bacterial species abundance) that were associated with the systemic immune system (see **Methods**). We hypothesized that CST IV bacteria or highly diverse CVMBs would be associated with plasma immune factors that are linked to adverse birth outcomes.

Shannon diversity was positively correlated with plasma MMP-2 (r = 0.214, p = 0.051) and CXCL13 (r = 0.353, p = 0.005) (**Fig. 7A-B**). When looking at individual taxa, *L. iners* was negatively correlated with sCD163 (r = -0.32, p =0.013 (**Fig. 7C**). In contrast, sTNFRSF1A concentrations were significantly higher in individuals with the CST III CVMBs vs those with CST IV CVMBs (**Fig. 7D**). However, in most cases, CST IV associated bacteria were positively correlated with immune markers of inflammation (**Fig. S3**). *L. crispatus,* a bacterium associated with low diversity in the CVMB, was negatively correlated with MMP-2 and CXCL13 and other biomarkers of inflammation (**Fig. S3**). In contrast, *L. gasseri,* also in low diverse CVMBs, was positively correlated with pro- and anti-inflammatory immune factors (**Fig. S3)**.

**Fig 7.**
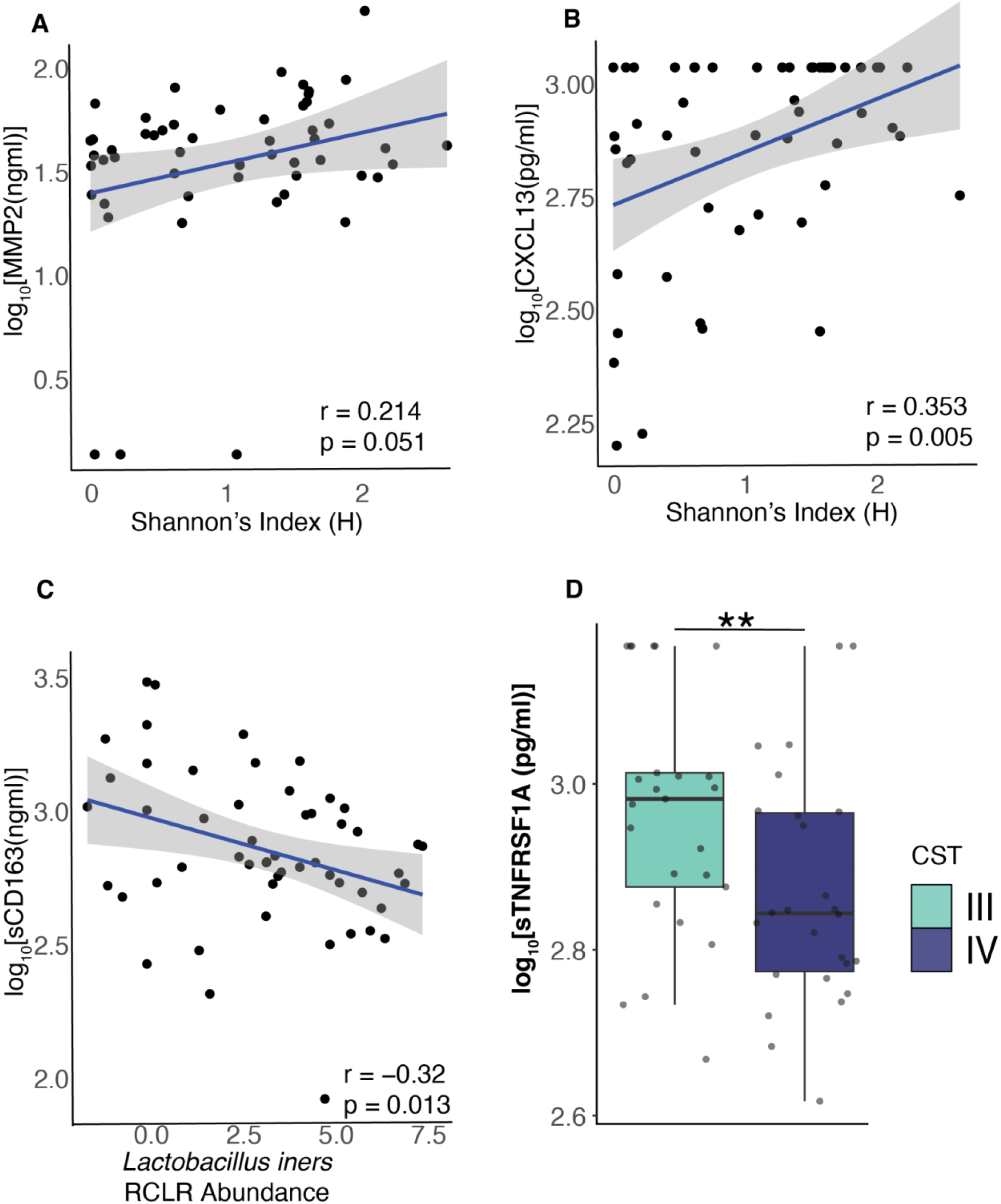
Correlations of CVMB metrics to plasma immune mediators A-B) Spearman correlation of MMP2 and CXCL13 to Shannon diversity **C)** sCD163 to *L. iners* abundance and **D)** Levels of sTNFRSF1A between CST III and IV (Wilcoxon Test p = 0.0188). Only statistically significant correlations (not corrected) are shown and only the top 3 abundant taxa are shown. Linear regression lines of A-C are shown in blue, and the shaded regions represent 95% confidence intervals. Full abbreviations of immume factors are in **Table S3**

Genital inflammation, linked to cervicovaginal dysbiosis, has been associated with adverse pregnancy and birth outcomes (21,101,102). Here, we investigate whether systemic inflammation is also linked to the CVMB. MMP-2 and CXCL13 were positively correlated with Shannon diversity. Abnormal concentrations and varying expression of MMP-2 during pregnancy have been implicated in adverse pregnancy outcomes like preeclampsia (103,104), recurrent spontaneous abortion (105), preterm birth (106) and gestational trophoblastic disease (107) and pregnancy outcomes may lead to adverse infant outcomes that impact infant mortality and development (108). The same is true for CXCL13 as it has been associated with placental inflammation (109) and preterm birth (110). Similarly, adverse pregnancy and birth outcomes, have been linked with highly diverse CVMBs, particularly those with CST IV communities (8,14,21,28). Altogether, this suggests that there is a complex interplay between the CVMB and systemic inflammation in PLWH-ART that impacts pregnancy and birth outcomes.

*L. iners* was negatively correlated with sCD163, an immune factor that has also been linked to preterm birth (111) and is a predictor of mortality in PLWH-ART (112,113) However, one study showed that sCD163 concentrations varied during pregnancy in PLWH-ART, declining in the first trimester and then remaining stable (114). All of this suggests that the role cervicovaginal bacteria have in modulating the systemic immune system in pregnant PLWH-ART — and therefore pregnancy and birth outcomes — is complex and may depend on the host’s immunological state. Indeed, the impact gut bacteria have on the systemic immune system might greatly confound the effect of cervicovaginal bacteria on the systemic immune system (115–117). Given that the local cervicovaginal immune profile is distinct from the systemic immune profile, cervicovaginal bacteria may have very little effect on the systemic immune profile (118). However, these associations between the plasma immune factors and maternal CVMB may be important for understanding the biological mechanisms underlying HIV-associated pregnancy and associated infant complications.

### Limitations, future opportunities and conclusions

The power of our study is limited by the lack of a pregnant PWoH group for comparison. This study was nested within a larger longitudinal study from the CQI-PMTCT study that was evaluating the long-term effects of ART usage in pregnant and chestfeeding people (119). As such, samples were only collected from pregnant PLWH-ART. Subsequent studies will have samples from pregnant PLWH-ART and PWoH and will also include larger sample sizes for better statistical inference. We did not present the impact of CVMB on birth outcomes in the CQI-PMTCT cohort as there were not enough samples in each adverse birth outcome to reach statistical significance.

Even acknowledging these limitations, our study revealed some important details about the CVMB of PLWH-ART in DRC and globally. We showed that the CVMB of pregnant PLWH-ART is largely CST III and IV both of which differ in their predicted functions. We found associations between the systemic immune system and the CVMB, providing direction for more studies to investigate this association as it might have implications on maternal and infant health. In a meta-analysis, we found that the HIV-associated CVMB and predicted function in pregnant PLWH-ART is distinct from PWoH, contrary to previous studies. Subsequent studies with larger cohorts in the same population in Kinshasa, DRC, will utilize shotgun metagenomic sequencing for genome-resolved analyses and better functional inference of bacterial and viral populations and their association with adverse birth outcomes in PLWH-ART. This will allow us to infer any strain level differences in the CVMB that may cause different host phenotypes, therefore ecosystem roles that are not detected with 16S rRNA gene sequencing. The CST framework would benefit to move on from taxonomy only based classification to microbial function and taxonomy-based classification that will give a better insight on the role of the CVMB on health and disease.

## Materials and methods

### Ethics approval and consent to participate

The Ohio State Institutional Review Board (number 2015H0440), Albert Einstein College of Medicine Institutional Review Board (number 2020-12018), and the University of Kinshasa School of Public Health Ethical Committee, and the University of Kinshasa School of Public Health Institutional Ethical Committee (ESP/CE/046/2018) granted approval for conducting this study and for protocol related specimen testing. All research was conducted in accordance with relevant guideline and regulations. Participants in the CQI-PMTCT study provided signed consent before undergoing study procedures

### Study cohort and sample collection

Participants in this analysis were enrolled in Kinshasa, Democratic Republic of Congo. This was a pilot study nested within the larger in the Continuous Quality Intervention: Preventing Mother to child transmission (CQI-PMTCT) study, a longitudinal study evaluation the long term outcomes of ART on pregnant people (NCT03048669) (120). From October 2020 to May 2021, we collected cervicovaginal swabs and peripheral blood from 82 participants at enrolment in their second or third trimesters who could access medical facilities during the SARS-CoV-2 pandemic. The 82 participants represent 40% of total participants who consented to provide cervicovaginal swabs. Clinicians collected cervicovaginal swabs and stored them in Puritan DNA/RNA shield (ZymoResearch, R1108) and transported to the central laboratory at Protestant University of Congo to be stored at - 80C until shipped to Columbus, OH on dry ice for processing. Plasma collected from the peripheral blood was processed immediately and stored at -80C until shipment to Columbus, OH for processing (**Fig. 1A**).

Demographic data including participant age, gestational age, ART regimen, gravidity, socioeconomic status, depression scores were collected at enrolment. The first principal component analysis (PCA) of the factors indicative of the participants socioeconomic status (SES) was used to categorize the SES status to quartiles with 0 as the lowest and 3 as the highest (121). Adverse birth outcomes including preterm birth, stillbirth and low birth weight were collected at delivery.

### Study selection, search strategy and criteria for meta-analysis

16S rRNA gene CVMB studies were selected by conducting a thorough and manually curated search of the Web of Science Core Collection of Thomson Reuters (WOS) and the European Nucleotide Archive (ENA) for studies published until December 2022. The search terms, hits and justification for exclusion or inclusion are shown in **Fig. S1** and information about the selected studies are in **Table S1**. The inclusion criteria were as follows: (i) pregnant healthy or living with HIV (iii) availability of raw data and (iii) Illumina paired-end short read amplicon sequencing of the V3-V4 hypervariable region. Only studies with sufficient information for comparison were included (i.e., metadata and minimally reproducible 16S rRNA gene sequence processing pipeline). Longitudinal studies were excluded from this analysis. Metadata from each study was obtained from NCBI, and/or via correspondence with the authors of the selected studies. Among these studies, three were from Europe –one from Italy (122) and two from Finland (123,124), one from the United States (125), and one from Uganda (126). Only the Ugandan study included samples from PLWH-ART (n = 5). In total, including this study, 1861 samples were analyzed. More details on the studies can be found in **Table S1.**

### DNA extraction, 16S rRNA gene sequencing and processing

Bacterial DNA from cervicovaginal swabs in the CQI-PMTCT cohort were extracted using the Qiagen AllPrep DNA/RNA Mini kit (Qiagen, 80204) following the manufacturer’s manual. These samples were low microbial biomass, hence, to reduce contamination, samples were extracted in a clean room and all materials were treated with 70% ethanol, RNAse and UV light for ∼15minutes before DNA extraction. Additionally, reagent-only controls were extracted in parallel with each DNA extraction batch. The PCR products of the reagent-only controls were examined on 1.2% gels, and no bands were detected.

Samples were sent to SeqCenter (https://www.seqcenter.com/) for amplification of the V3-V4 hypervariable region of the 16S rRNA gene using the 341F (5’- CCTACGGGDGGCWGCAG-3’) and 806R (5’-GACTACNVGGGTMTCTAATCC-3’) primers using the Zymo Research’s Quick-16S kit. The V3-V4 region was chosen as it has been shown to identify the most taxa in cervicovaginal samples (127). The PCR products were sequenced on an Illumina NextSeq2000 to generate 2x301bp paired-end reads.

16S rRNA gene reads were processed following the DADA2 SOP (128). Sequence quality was assessed using FASTQC v0.11.5 and any adapters or primer sequences were trimmed using Trimmomatic v0.36 (129,130). Generally, sequences were trimmed at the first base with a quality score of Q<25. Forward and reverse reads were merged using the mergePairs function with default parameters (minOverlap = 12, maxMismatch = 0), and chimeras removed using removeBimeraDenovo with default parameters. Taxonomy was assigned using the Greengenes2 database (131). Sequences were filtered to remove sequences of mitochondria and chloroplast, samples with less that 100 reads and taxa with less than 10 reads, less than 0.5% abundance across all samples and not present more than 2 times in 10% of samples were removed. The resultant ASV table was agglomerated to species-level and robust centered log-ratio (RCLR) transformation was performed on the table using the decostand function on the vegan package v2.6-6. This transformation was to mitigate the inherent negative correlation bias in compositional data (132,133) The transformed table was used in all subsequent analyses.

For the five (5) selected studies for the meta-analysis, sequences were processed on a per-study basis as described above. Reads were filtered based on read quality for each study. Per study processing parameters are shown in **Table S2.** Generated ASV tables per study were combined and taxonomy was assigned on the combined table (34,871 ASVs and 1861 samples). The aggregated table was further filtered by removing ASVs not assigned to any taxonomic level. To reduce data complexity (134), bacterial ASVs were filtered if observed at frequencies of 0.0005% study-wide and observed in only 2 samples. Samples with fewer than 2000 reads or with excess zeroes were removed using the PreFL function from the PLSDAbatch package v1.0 (135). After filtering, ASVs were agglomerated at the species level resulting in an abundance table with 74 species-level taxa and 664 samples. A centered log-ratio (CLR) transformation was performed on the abundance table. Before the CLR transformation, zeros were replaced with a constant value smaller than the detection limit (e.g., 65% of the detection limit) (136,137).

PICRUSt2 version v2.4.2 (79) was used to predict functional composition from the 16S rRNA gene sequences. Data were normalized based on the abundance of 16S rRNA gene copy numbers. Metagenome predictions were used to estimate KEGG pathways and any unassigned KEGG IDs were mapped using KEGGREST v3.19 (138).

### Quantification of plasma cytokine, chemokine and soluble factors (immune factors)

Levels of cytokines, chemokines and soluble factors were measured from maternal plasma were quantified using the LEGENDplex COVID-19 Cytokine Storm Panel 1 & 2 (BioLegend, 741095), LEGENDplex Human Proinflammatory Chemokine Panel 1 (BioLegend, 740984) and LEGENDplex Vascular Inflammation panel 1 (BioLegend, 740551). Additional details related to measurement of immune factors are described in Supplemental Methods.

### Statistical analyses

For the CQI-PMTCT study, statistical analyses were as follows. Community state types (CST) were assigned to each sample using VALENCIA, which assigns CSTs to each samples based on their similarity to a reference centroid (139). Alpha diversity indices (Chao1, Simpson, and Shannon) were estimated using an asymptotic statistical approach (140,141). Differences in alpha diversity were tested using the Kruskal-Wallis test, followed by Wilcoxon pairwise comparison tests if significant.

Beta diversity analyses were computed using principal component analysis (PCA) on Robust Aitchison distances. Significance of PCAs used ANOSIM from the vegan package v2.6-10. To identify associations between CVMB metrics (alpha diversity and bacterial abundance) and the log transformed plasma cytokine, chemokine and soluble factors concentrations, linear regression models were computed with the with Spearman’s rank correlation. Adjustments for multiple hypothesis testing were not performed due to this being an exploratory study unless as a sensitivity analysis for our main findings (142)

For the meta-analysis, data were analyzed as following. CSTs were assigned as described above using VALENCIA. Batch effects were accounted for using Partial Least Square Discriminant Analysis (PLSDA-batch) after evaluation of several batch removal methods (see Supplemental Methods for more details). Alpha diversity indices and beta diversity on Aitchison distances were computed as above.

To identify differentially abundant species and predicted functions, three statistical methods were used: LinDA from the MicrobiomeStat package v.1.1 (143,144) , ANCOM-BC v.2.2.1(145) and ALDEx2 v.1.32.0 (146) with the formula “abundance ∼ health_status + study”. A species was considered differentially abundant if it met the following criteria: p-adjusted value < 0.05 in LinDA; q-value < 0.05 in ANCOM-BC; or effect size > 0.5 in ALDEx2. Additional details on differential abundance analysis are described in Supplemental Methods. The STORMS checklist (147) (**Table S5**) is included for standardization and reproducibility of data.

## Data availability

The datasets supporting the conclusions of this article are available in the Sequence Read Archive (SRA) under BioProject PRJNA1234224 and scripts and data used to analyses all the data are available on https://github.com/SulliVagOSU/16S_meta_analysis.

## Acknowledgments

This work was funded by the National Institute of Child Health and Human Development (NICHD) under the award numbers R01HD087993 and R01HD105526. The funders had no role in study design, data collection and analysis, interpretation and preparation of this manuscript. The authors declare no competing interests.

We thank all the people who participated and contributed to this study. We also thank the contribution of the CQI-PMTCT study team, the staff of the participating clinics and the provincial and national health authorities. We would also like to thank members of the Sullivan and Kwiek labs for their support and critical feedback. Bioinformatics analysis was supported in part by the Ohio Supercomputer Center. Figure 1A and 1B were created in BioRender.

## Notes

### Competing Interest Statement

The authors have declared no competing interest.

### Summary of Updates

Added an author (Ann C. Gregory).

