## Supplemental data for "The cervicovaginal microbiome of pregnant people living with HIV on antiretroviral therapy in the Democratic Republic of Congo: A Pilot Study and Global Meta-analysis"

Correpsonding authors:

**Matthew B. Sullivan**

**Jesse J. Kwiek**

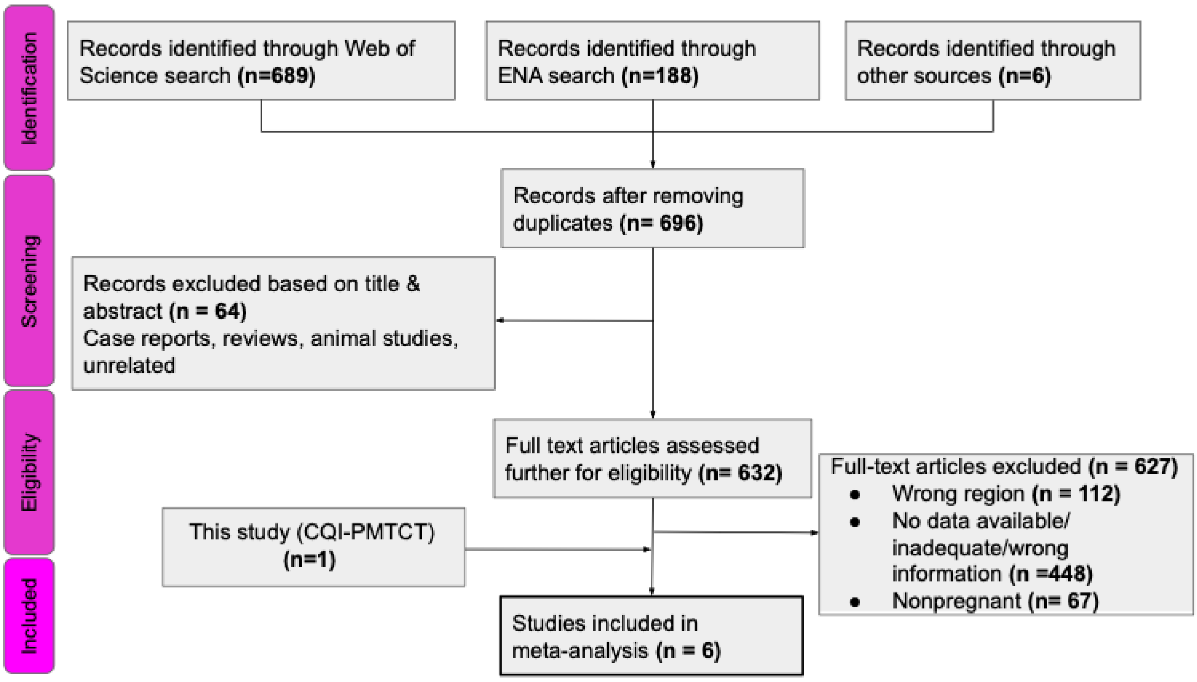


**Supplemental Figure 1:** PRISMA flowchart on selection of studies for meta-analysis. The search terms for ENA were: “vaginal microbiome” and for WOS: “(ALL= (country name OR nationality) AND TS= (vaginal OR vagina OR vagin*) AND TS= (microbiota* OR microflora* OR bacteria* OR microbiome* OR flora* OR bacterial* OR bacteria* OR microorganism OR dysbiosis) AND TS=(16S) AND TS= (woman OR women))”

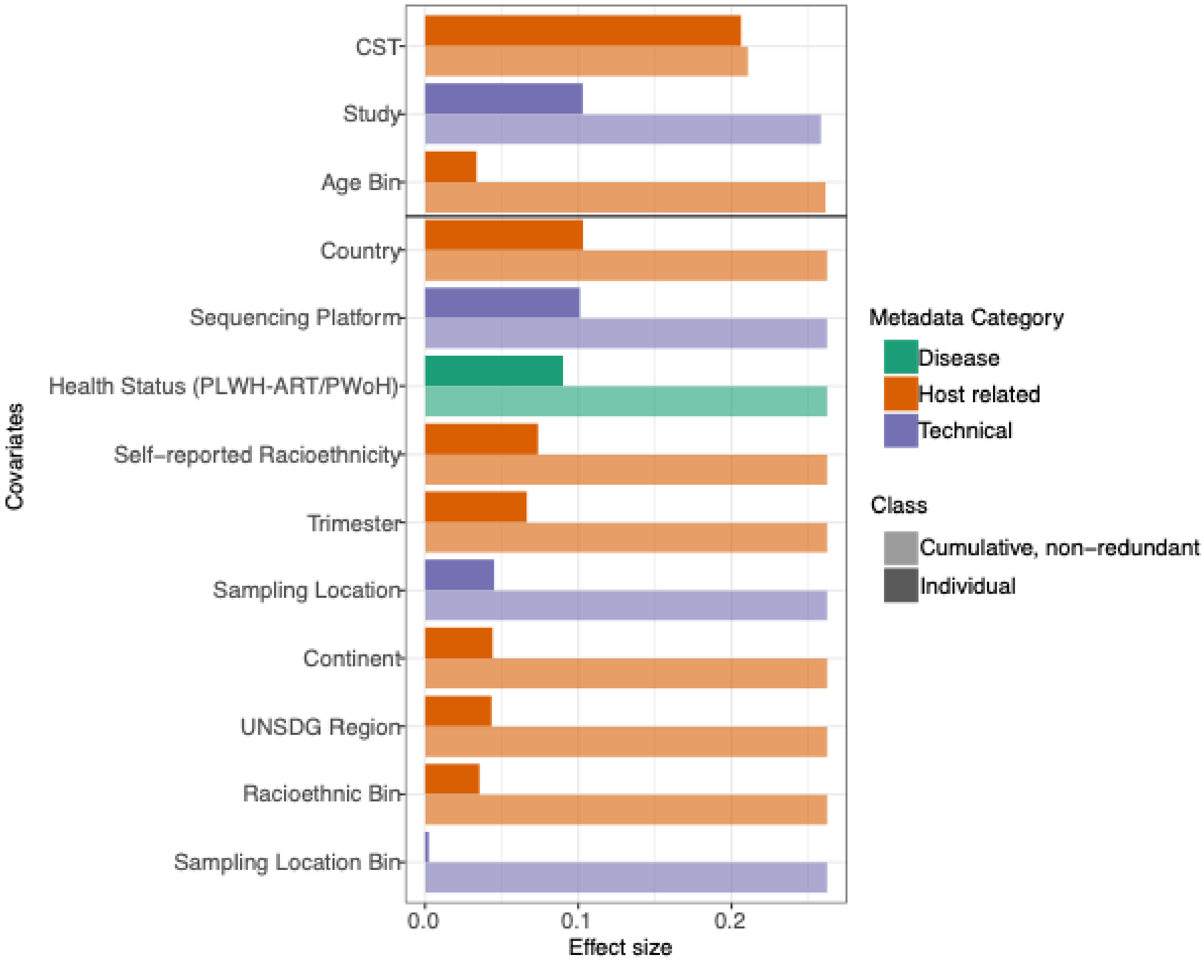


**Supplemental Figure 2:** Covariates explaining microbial variation in the CVMB of the 6 studies included in the meta-analysis (n=625, dbRDA). Only significant covariates shown (BH-corrected $p < 0.05$). Dark colors indicate the individual variance explained by each covariate and lighter colors show the cumulative, non-redundant variance explained by the covariates (n =625, stepwise dbRDA). The black line represents only the covariates that were significant in the cumulative, non-redundant analysis.

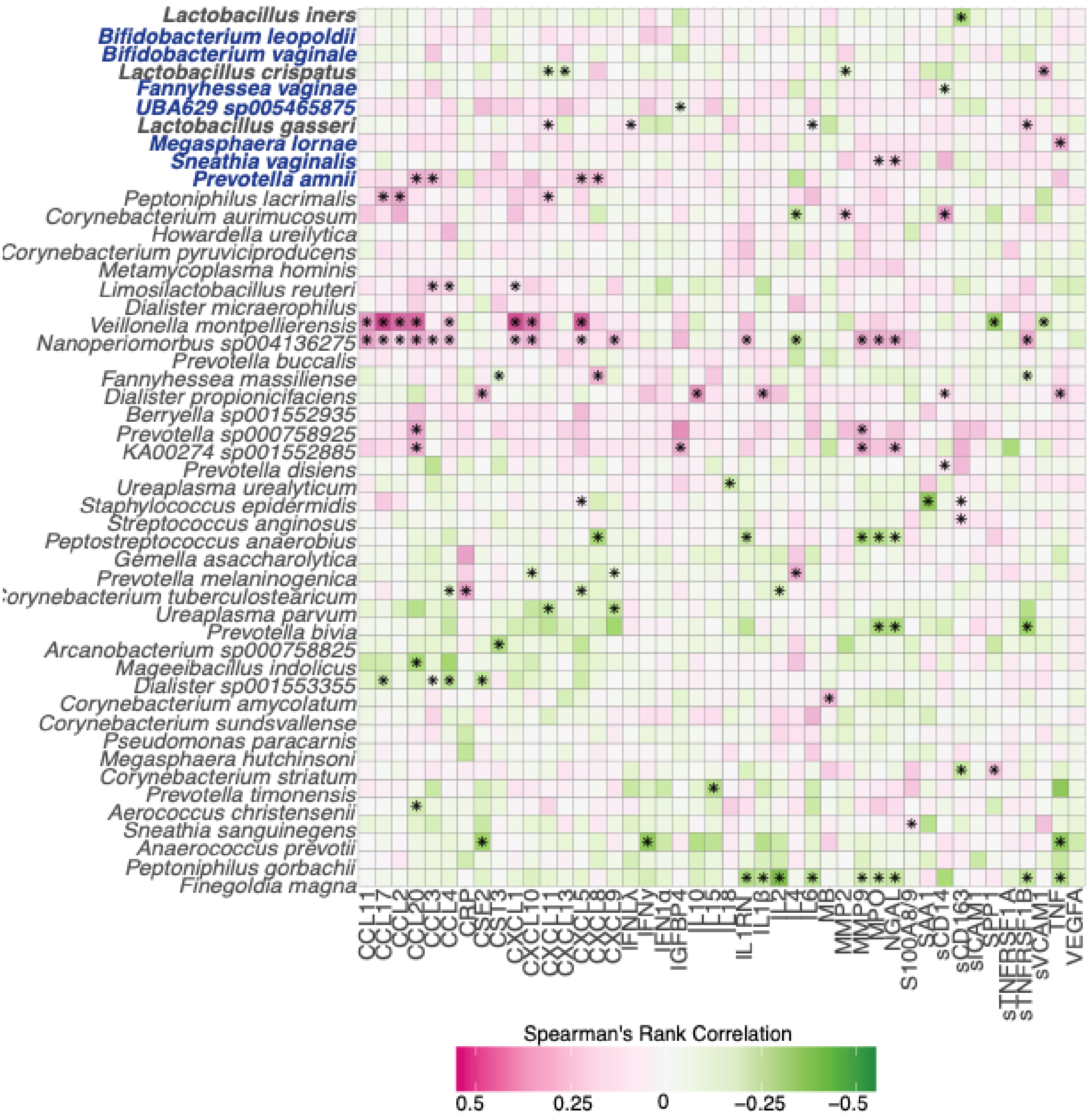


**Supplemental Figure 3:** Heatmap showing spearman correlations between taxa and log10-normalized concentrations of immune factors. The top 10 most abundant taxa are bolded, and CST IV associated bacteria are highlighted in blue. Taxa are ordered by their overall abundance. Asterisk indicates spearman's $p < 0.05$, not adjusted. Filled circle indicates FDR corrected $p < 0.05$.

**Supplemental Methods**

***Quantification of plasma cytokine, chemokine and soluble factors (immune factors).***

All immune factor names, abbreviations and measurement units (ng/ml or pg/ml) can be found in **Supplemental Table 3**. Names and abbreviations were curated from uniport.org, *homo sapiens*. Names and abbreviations were curated from uniport.org, *homo sapiens,* following the International Protein Nomenclature Guidelines (1). Thirteen chemokines were simultaneously measured in plasma using LEGENDplex Human Proinflammatory Chemokine Panel 1 (BioLegend, 740984) (interleukin-8 (IL-8), C-X-C motif chemokine 10 (CXCL10), eotaxin (CCL11), C-C motif chemokine 17 (CCL17), CCL2, CCL5, CCL3, CXCL9, CXCL5, CCL20, CXCL1, CXCL11, CCL4) according to the manufacturer's filter plate protocol. Similarly, twelve cytokines were measured in plasma using the LEGENDplex COVID-19 Cytokine Storm Panel 1 & 2 (BioLegend, 741095) (IL-6, interferon alpha-2 (IFN-α-2), IL-2, IFN-ɣ, IL-1RN, tumor necrosis factor (TNF- α), IL-10, granulocyte-macrophage colony-stimulating factor (GM-CSF), IL-1β, vascular endothelial growth factor a, long form (L-VEGF), IL-18, IL-15). Thirteen cytokines and soluble factors were measured in plasma using the LEGENDplex Vascular Inflammation panel 1 (BioLegend, 740551) (myoglobin (MB), protein S100-A8/protein S100-A9 (S100A8/S100A9), neutrophil gelatinase-associated lipocalin (NGAL) , c-reactive protein (CRP), 72 kDa type IV collagenase (MMP-2), osteopontin (SPP1), myeloperoxidase (MPO), serum amyloid A-1 protein (SAA), insulin-like growth factor-binding protein 4 (IGFBP4), soluble intercellular adhesion molecule 1 (sICAM1), soluble vascular cell adhesion protein 1 (sVCAM1), matrix metalloproteinase-9 (MMP-9), cystatin-C (CST3). Data were collected on the MACS Quant 10 Flow cytometer (Miltenyi

Biotech) and using MacsQuant Software (Miltenyi Biotech). Gates were set around beads to exclude debris, and number of events collected was 300 per analyte multiplied by 1.1. Gates around individual bead sets and the corresponding gates around individual analytes were adjusted post collection in the LEGENDplex online Qognit software based on standards; gates were then applied to all standards and samples for a particular plate and five-parameter logistic standard curves were generated for all. Concentrations were interpolated from these curves using the LEGENDplex Qognit software (BioLegend, Qognit Inc.).

Seven immune factors were individually measured by duoset ELISA (R&D Systems): CXCL13 (DY801), IL-4 (DY204), IFN-λ-1(DY7246), soluble monocyte differentiation antigen CD14 (sCD14, DY383), soluble scavenger receptor cysteine-rich type 1 protein M130 (sCD163, DY1607), soluble tumor necrosis factor receptor superfamily member 1A (sTNFRSF1A, DY225), sTNFRSF1B (DY726) according to the manufacturer's instructions with minor modifications. First half-well plates were used (Greiner, 675061), plates were washed using the CAPP wash 12 (Pipette.com, W-12) attached to a carboy that was washed daily, and plates were developed using TMB substrate Plus liquid (VWR, 97063-666). Absorbances were read on a SpectraMax i3x (Molecular Devices).

For all immune factor assays performed there were three samples that were on all plates to serve as quality control to ensure similar assay performance across plates and days. See **Supplemental Table 4** for dilution factors and upper and lower limits of detection (LoD). Values below the limit of detection or below the bottom standard were increased to the bottom standard, unless the dilution factor was >2, then the bottom standard was multiplied by the dilution factor. Values above the limit of detection were replaced by the top standard multiplied by the dilution factor multiplied by 1.1 (**Supplemental Table 4**).

***Statistics***

To avoid batch effects, removeBatchEffect (rBE from the limma package) (2), ComBat from the sva package in R (3), Partial Least Square Discriminant Analysis (PLSDA-batch), sparse PLSDA (sPLSDA-batch), weighted PLSDA (wPLSDA-batch), sparse weighted PLSDA (wPLSDA-batch), from the PLSDAbatch package, and batch-mean centering transformation (BMC). To evaluate batch adjustment correction the following methods were used: principal component analysis (PCA); heatmap and cluster analysis using tidyHeatmap v. 1.8.1 (4); alignment score (PLSDAbatch package); and partial redundancy analysis (pRDA) using vegan v. 2.6-4 (5).

To identify differentially abundant species and predicted functions, three statistical methods were used: LinDA from the MicrobiomeStat package v.1.1 (6,7) , ANCOM-BC v.2.2.1(8) and ALDEx2 v.1.32.0 (9) with the formula "abundance ~ health_status + study". A species was considered differentially abundant if it met the following criteria: p-adjusted value < 0.05 in LinDA; q-value < 0.05 in ANCOM-BC; or effect size > 0.5 in ALDEx2. A score was computed to indicate the number of methods that identified a species as differentially abundant. For example, if a species was identified by both LinDA (p-adjusted value < 0.05) and ALDEx2 (effect size > 0.5), it received a score of 2. This combination of methods aims to overcome drawbacks such as low power and high false discovery rate (FDR) in differential abundance methods (10). The input was the count table without CLR transformation. In LinDA, a heuristic imputation method was used, with imputed values proportional to library sizes. ANCOM-

BC and ALDEx2 included their own transformations and bias corrections. The analysis was conducted using default arguments for all methods.
